# Transcriptomics of improved fruit retention by hexanal in ‘Honeycrisp’ reveals hormonal crosstalk and reduced cell-wall remodelling in the fruit-abscission zone

**DOI:** 10.1101/2021.05.19.444856

**Authors:** Karthika Sriskantharajah, Walid El Kayal, Davoud Torkamaneh, Murali Mohan Ayyanath, Praveen Saxena, Alan J Sullivan, Gopinadhan Paliyath, Jayasankar Subramanian

**Author notes:** Author for correspondence: Jayasankar Subramanian *Email:.

## Abstract

Apples (*Malus domestica* Borkh) are prone to pre-harvest fruit drop which is more pronounced in ‘Honeycrisp’. Using a transcriptomic approach, we analyzed the molecular mechanisms of fruit retention in ‘Honeycrisp’. A total of 726 differentially expressed genes (DEGs) were identified in the abscission zone of hexanal-treated and untreated fruit (FAZ). Hexanal down-regulated the genes involved in ethylene biosynthesis, such as S-adenosylmethionine synthase (*SAM2*) and 1-aminocyclopropane-1carboxylic acid oxidases (*ACO3*, *ACO4* and *ACO4*-*like*). Genes related to ABA biosynthesis (*FDPS* and *CLE25*) were also down-regulated. On the contrary, gibberellic acid (GA) biosynthesis genes, gibberellin 20 oxidase1-like (*GA20OX*-*like*) and ent-kaurene oxidase (*KO*) were up-regulated. Further, hexanal down-regulated the expression of genes related to cell-wall remodelling enzymes such as polygalacturonase (*PG1*), glucanases (endo-β-1,4-glucanase; EG) and expansins (*EXPA1*-*like, EXPA6, EXPA8, EXPA10*-*like, EXPA16*-*like*). Hexanal also reduced ethylene, and abscisic acid (ABA) production at commercial harvest stage. Hexanal reduced ethylene production in fruits and thus reduced the sensitivity of FAZ cells to ethylene and ABA. Simultaneously, hexanal maintained the cell-wall integrity of FAZ cells by regulating genes involved in cell-wall modifications. Our findings show that fruit abscission is delayed by hexanal, by down regulating ABA through an ethylene-dependent mechanism.

**Highlight:** Hexanal, a naturally occurring plant compound, increased fruit retention in apples by decreasing ethylene and ABA production and maintaining the cell-wall integrity in the fruit abscission zone.

## Introduction

Apple (*Malus domestica* Borkh.) is one of the most widely cultivated temperate fruits and ranks third in global fruit production. Most apple trees tend to shed fruits just before the harvest, often referred to as pre-harvest fruit drop (PFD), which renders a huge economic loss to growers. The PFD usually begins 3-4 weeks before the anticipated harvest and causes yield losses up to 30% at the beginning of the harvest (Robinson, 2011; Arseneault and Cline, 2016). The severity of the PFD is cultivar specific and influenced by the orchard and climatic factors. Moreover, this issue is exasperated when the fruits are left on the tree for better colour development, something that has a huge consumer appeal. ‘Honeycrisp’, a premium apple cultivar, is categorized as more prone to PFD (Irish-Brown *et al*., 2011), which causes yield losses of almost 50% in some years (Arseneault and Cline, 2017).

Pre-harvest fruit drop is a consequence of abscission, whereby cell separation occurs rather pre-maturely at the constriction region of the pedicel resulting in fruit drop (Addicott, 1982; Taylor and Whitelaw, 2001). The PFD control measures in apples have largely relied upon the use of plant growth regulators (PGRs) and are often cultivar-specific. Moreover, abscission is an irreversible physiological process, thus warranting suitable technologies and methods to improve fruit retention and overcome orchard management challenges. Application of hexanal as a formulation at the pre-harvest stage has shown promising results in improving fruit retention in several fruits, including apple (DeBrouwer *et al*., 2020), raspberry (EI Kayal *et al*., 2017), mango (Anusuya *et al*., 2016), and orange (Samwel *et al*., 2020). Hexanal application also extends the shelf life of several horticultural commodities through inhibiting the membrane degradation enzyme, phospholipase-D (Paliyath *et al*., 2003; Paliyath and Subramanian, 2008). Previous studies on fruit abscission have reported that the cell-separation within the FAZ is the result of a cascade of physiological events due to the coordinated expression PGR related genes resulting in abrupt changes in endogenous levels of plant hormones (Addicott, 1982; Estornell *et al*., 2013; Kumar *et al*., 2013).

Plant hormone ethylene is the key regulator of abscission. Application of ethylene inhibitors such as aminoethoxyvinylglycine (AVG) and 1-methylcyclopropene (1-MCP) reduced the PFD in ‘Bisbee Delicious’ apples (Yuan and Li, 2008). On the contrary, ethylene releasing compound, ethephon, promoted fruit abscission, indicating that both ethylene biosynthesis and signalling pathways are involved in fruit abscission. Fruit ethylene production and fruit softening increased rapidly during fruit ripening of PFD-prone cultivar ‘Golden Delicious’ than non-prone cultivar ‘Fuji’ (Li *et al*., 2010). Moreover, transcript levels of ethylene biosynthesis genes *MdACS5A*, *MdACO1*, and receptor genes *MdETR2* and *MdERS2* increased in the FAZ of ‘Golden Delicious’ (Li *et al*., 2010). Similarly, early induction of fruit abscission in melon was associated with up-regulated expression of *SAMS*, *ACS, ACO* and *ETRs* in the FAZ (Corbacho *et al*., 2013). However, efforts to suppress the expression of ethylene related genes using PGRs effectively worked only in combined applications (Yuan and Carbaugh, 2007; Greene, 2009; Robinson *et al*., 2010). Previous studies have observed that hexanal decreased ethylene production in the ripening fruits (Jincy *et al*., 2018; DeBrouwer *et al*., 2020) and significantly down-regulated the expression of the *ACS* gene (Tiwari and Paliyath, 2011). However, in spite of its promise in improving fruit retention, there is no information on how hexanal regulates the fruit abscission.

Plant hormones, particularly abscisic acid (ABA), auxin and gibberellins (GA), also play substantial roles in fruit abscission. ABA levels generally increase towards fruit maturity and ripening and contribute to senescence and seed dormancy (McAtee *et al*., 2013). Although a high level of ABA in the AZ, prior to abscission, has been reported in several species, the direct involvement of ABA in the abscission process remains unclear. It has been suggested that ABA appears to act as a modulator of ACC levels, and therefore stimulates ethylene biosynthesis, leading to increased abscission (Guinn, 1982; Wilmowicz *et al*., 2016). Exogenous application of ABA contributed to PFD in ‘Golden Delicious’ apples (Masia *et al*., 1998). Abscission was delayed in the ethylene-JA-ABA deficient triple mutant in Arabidopsis and showed an association between abscission and ABA levels (Ogawa *et al*., 2009). Contrary to ABA, auxin flux across the FAZ reduced the ethylene sensitivity of AZ and subsequently increased fruit retention (Taylor and Whitelaw, 2001; Estronell *et al*., 2013). Moreover, maintenance of polar auxin transport contributed to improving fruit retention in sweet cherries (Blanusa *et al*., 2005).

Cell wall breakdown and cell separation are required within the FAZ for the fruit to abscise from the tree. Cell wall hydrolysis enzymes such as polygalacturonase (PG), cellulase (EG) pectate lyase and cell wall loosening enzymes expansins promote fruit detachment at the AZ (Bonghi *et al*., 2000; Li and Yuan, 2008). Ethylene is strongly correlated with the activity of these hydrolases and the expression of genes including *MdPG2* and *MdEG1* in the FAZ (Li *et al*., 2010). These findings suggest that ethylene plays a regulatory role in fruit abscission and can accelerate the abscission process. Ethylene also appeared to interact with other plant hormones in fruit drop, particularly with ABA (Masia *et al*., 1998) and auxin (Blanusa *et al*., 2005).

We previously reported that the pre-harvest hexanal spray improved post-harvest storage attributes of ‘Honeycrisp’ apples (DeBrouwer *et al*., 2020). Based on those studies, we hypothesize that hexanal improves fruit retention either by regulating the abscission through an ethylene dependent mechanism or directly manipulating the ABA/GA mechanism. To test this hypothesis, we studied the mechanism of action of hexanal in improving fruit retention in ‘Honeycrisp’ using multidisciplinary approaches including physiological, biochemical, and transcriptomic characterization. The differentially expressed genes (DEGs) due to the application of hexanal in the FAZ were identified via RNA-seq analysis. The functional profiling of the DEGs were studied through gene ontology annotation and enrichment analysis. Plant hormones present in the FAZ were quantified using reverse-phase ultra-performance liquid chromatography-mass spectrometry.

## Materials and methods

### Trial location and pre-harvest treatment

Field trials were conducted at two commercial apple orchards located within the Niagara region of Ontario, Canada, (Site A; 43°08′53.7” N, 79°29′50.2” W and Site B; 43°11′00.1” N 79°34′44.4” W). Both sites had ‘Honeycrisp’ trees grafted onto M9 rootstocks supported by a trellis system. The trees at Site A were eight years of age, and the trees at Site B were nine years of age. Hexanal formulation (HF) was prepared as described earlier (EI Kayal *et al*. 2017; Kumar *et al*. 2018), containing hexanal at a concentration of 0.02 % (v/v) in the final spray. ‘Honeycrisp’ trees were subjected to two pre-harvest sprays of HF approximately 30 and 15 days before the commercial harvest. Trees sprayed with water served as control. Buffer rows were maintained between the treatments to avoid spray contamination. A total of 48 trees from each site were used for the study.

### Plant material, RNA-isolation, and library preparation

Fruit stalk samples on the constriction region between pedicel and spur of about 3-4 mm (FAZ) were harvested on the commercial harvest day from both sites (Fig. 1). Freshly excised FAZ samples were flash frozen in liquid nitrogen at the field and stored at −80 °C for RNA isolation. Total RNA was extracted from FAZ tissues using an RNA isolation kit (Norgen Biotek, ON, Canada). RNA quality was verified and quantified using Nanodrop^TM^ (2000/2000c Spectrophotometers, Thermo Fisher Scientific, USA). One microgram of mRNA was used as a template for first-strand cDNA synthesis using NEBNext® Poly(A) kit (NEB #E7490, New England Biolabs, Inc.) and NEBNext® ultra™ II directional RNA library prep kit for Illumina (NEB #E7760, New England Biolabs, Inc.). Paired-end sequencing (75 bp) was performed for four samples using NextSeq 500/550-mid output kit v2.5 (2 × 75 cycles) on an Illumina NextSeq500 sequencer (Norgen Biotek, ON, Canada).

**Fig. 1:**
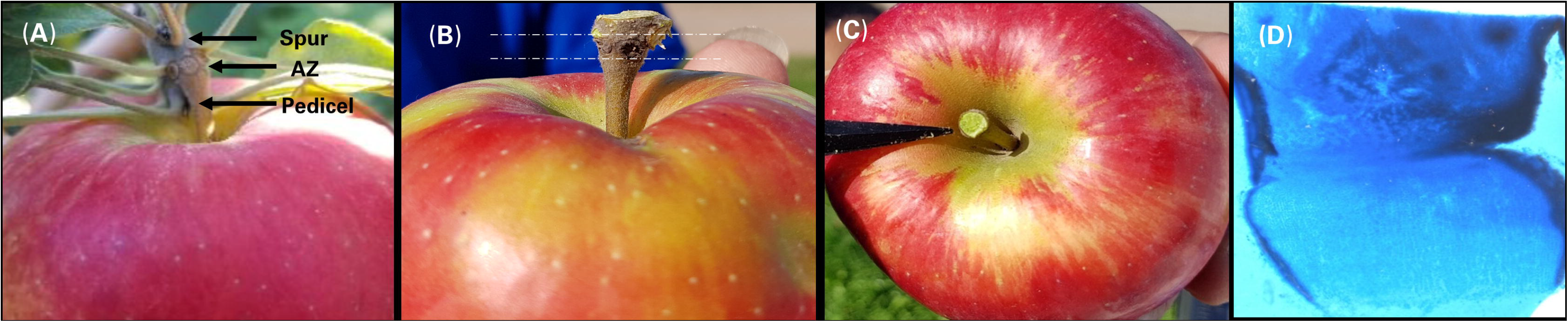
Anatomical observation of the FAZ of ‘Honeycrisp’ apple. (A), (B) photographs show the location of the abscission zone in the fruit, (C) image of the excised AZ plane without any mechanical damages and (D) microscopic view of the AZ region. The AZ was stained using lactophenol cotton blue. The microscopic view of the AZ looks like a funnel shape.

### Trimming, assembly, and annotation of paired-end sequenced reads

The quality of raw sequences was measured with FastQC (Andrews, 2010) and were trimmed by Trimmomatic (v.0.36) (Bolger *et al*., 2014). Then adapter sequences used for library preparation were removed with Cutadapt using sequencing by oligonucleotide ligation and detection colour space algorithm (SOLiD, Martin, 2011). Trimmed reads were assembled with STAR aligner (v2.1.3) with default parameters (Dobin *et al*., 2012). Apple genome project GDDH13, version1.1 was used as a reference genome (Daccord *et al*., 2017). FeatureCounts program was used to assign sequence reads to each gene in the samples corresponding to the reference genome. A table of count reads was created with rows corresponding to genes and columns to samples (Liao *et al*., 2014).

### Differentially expressed gene analysis

EdgeR (Robinson *et al*., 2010), an R Bioconductor package deposited in the DEBrowser (v1.16.1) (Kucukural *et al*. 2019), was used to analyze the differentially expressed genes (DEGs) between hexanal treated and control samples at *p* ≤ 0.05 and gene expression fold change ≥ 2. The EdgeR models count data using an over-dispersed Poisson model and an empirical Bayes procedure to moderate the degree of overdispersion across genes. Table of count reads of four samples was fed to the EdgeR program. DEGs were analyzed through data assessment, normalization and DEG detection using EdgeR models. The EgdeR modelled data to negative binomial (NB) distribution, Y_gs_∼NB(X_s_Z_gs_, □_g_) for gene g and sample s. Here X_s_ is the library size, □_g_ is the dispersion, and Z_gs_ is the relative abundance of gene g into which sample s belongs. The NB distribution reduces to Poisson when □_g_ = 0 (Robinson *et al*., 2010).

### Enrichment analyses

Gene ontology (GO) and functional pathway enrichment analyses were performed in ShinyGO (v0.61), based on hypergeometric distribution followed by Benjamini-Hochberg correction with a false discovery rate (FDR) at *p* ≤ 0.05 (Ge *et al*., 2019). Three different gene ontologies, such as biological process (BP), molecular function (MF) and cellular components (CC), were analyzed separately. The relationship between enriched functional pathways was visualized using an interactive plot. A hierarchical clustering tree was also used to summarize the correlation among the enriched pathways. *Arabidopsis thaliana* (TAIR10) ortholog genes for the identified *Malus domestica* DEGs (726) were retrieved from the phytozome database (v13.1.6) and were used for the enrichment analysis (Balan *et al*., 2018). Query-based gene information of all detected DEGs was obtained using the phytozome database for the *Malus* domestica (GDDH13, v1.1).

### Quantitative RT-PCR

Quantitative reverse transcription PCR was conducted for eight genes chosen to represent ethylene biosynthesis and signalling pathway and cell-wall modification. Gene-specific primers were designed using Primer3Plus software (Untergasser *et al*., 2012). Two micrograms of total RNA extracted from FAZ were reverse transcribed with Superscript II reverse transcriptase (Invitrogen, Canada). qPCR reactions were performed in 10 μL, containing 5 μL SYBR^®^ Green Supermixes (Invitrogen, Canada), 50 ng of cDNA and 2.5 μL of 400 nM of each primer (Table **S1**). Four biological and three technical replicates for each gene were analyzed using a CFX96 Real-Time PCR detection system (BioRad, Mississauga, ON, Canada). *Malus domestica Actin* (*MdACT*) and *Histone-3* (*MdHIS-3*) genes were used as reference genes to normalize the gene expression of a target gene. The gene expression was quantified using the 2^−ΔΔCt^ method (Livak & Schmittgen, 2001).

### Fruit retention and fruit quality measurements

Four trees with uniform growth, similar maturity, comparable fruit count, and similar location for wind direction were marked for fruit retention study. Fruit retention (FR) was monitored on a biweekly basis from the commercial harvesting in September to before the first frost in November and expressed as a percentage using the following formula:

Fruit Retention % = 100 − [(initial fruit count − final fruit count) / initial fruit count) × 100]

Ten randomly selected, similar-sized fruits at commercial harvesting (0 days) and end of the fruit retention study (49^th^ day) were used for the quality measurements. Fresh weight (g) was measured using an electronic balance. Two firmness readings (N) were taken using a handheld penetrometer with an 11 mm diameter tip (Effegi pressure tester, Facchini 48011, Alfonsine, Italy) on the opposite sides of each fruit. Two vertical slices from each side of the apples were freshly juiced, and TSS (^o^Brix) readings were measured using a prism refractometer (Fisher Scientific, Mississauga, Canada).

### Plant hormone measurement

Eight randomly selected apples were used for the ethylene measurement. Fruits were weighed and placed in 2 L glass bottles. Bottles were sealed for an hour with a lid containing a rubber port where a syringe was used to collect 1 mL of headspace gas after gently shaking the bottles to mix up the air inside. The gas sample was immediately injected into an SRI-8610c gas chromatograph equipped with a 0.5 mL sample loop, and ethylene was detected using a flame ionization detector (Varian Inc., Mississauga, ON).

Plant hormones present in the FAZ were extracted using the methanol double extraction method. Briefly, 25 mg of freeze-dried, powdered FAZ samples were extracted with a solvent (methanol: formic acid: milli-Q H_2_O = 15:1:4), and the homogenate was kept at −20 °C for an hour. The supernatant was then collected through centrifugation (15 min, 14,000 rpm). The pellet was re-extracted using the same protocol, and the supernatants were pooled. The pooled supernatant was then evaporated to dryness using nitrogen gas in a fume hood. The dried samples were reconstituted using a buffer solution (0.1% formic acid: acetonitrile = 97:3) and was then filtered through a 0.45 μm centrifuge filter (Millipore; 1 min, 13,000 rpm). The supernatant was then transferred to a 96-well collection plate. Metabolites were separated by reverse-phase ultra-performance liquid chromatography (UPLC) system with detection using an Aquity QDa single quadruple mass spectrometer (MS) controlled by Empower 3 (Waters, Canada) by injecting a 5 μL aliquot of sample onto an Acquity BEH Column (2.1 × 50 mm, i.d. 2.1 mm, 1.7 μm). Metabolite peaks were monitored in single ion recording mode and quantified using a standard curve (Erland *et al*., 2017).

### Statistical analysis

Fruit retention, quality and hormone data were analyzed using general linear mixed models (proc GLIMMIX) in SAS v9.4 (SAS Institute, Raleigh, NC). Variance of fixed effects such as location and treatment were partitioned from random effects, which include replication. Shapiro-Wilk normality tests and studentized residual plots were used to test error assumptions of variance analysis including random, homogenous, and normal distribution of error. Means were calculated using the LSMEANS statement, and significant differences between the treatments were determined by the Tukey-Kramer test with α = 0.05 and are mentioned in each figure or table.

## Results

### Fruit retention and fruit quality as a result of hexanal spray

Hexanal application significantly and consistently increased the fruit retention in the treated trees compared to control (Fig. 2). All trees showed a continuous decline in fruit retention throughout the postharvest, but the rate of decline was significantly slower in the treated trees than in control. At the end of the fruit retention study period of 49 days, treated trees retained three-quarter (74%) of the total fruits while control trees retained less than half (46%) of the total fruits. In apples, retaining fruits for a longer duration facilitates natural ripening and colour development on the trees and improves fruit quality. Hexanal treated fruits had significantly higher firmness than control fruits. The firmness rapidly declined in the control fruits while the decline was quite slow in the hexanal treated fruits. However, other quality parameters did not vary significantly (Table 1), indicating that hexanal does not alter any other fruit quality characteristics.

**Table 1:**
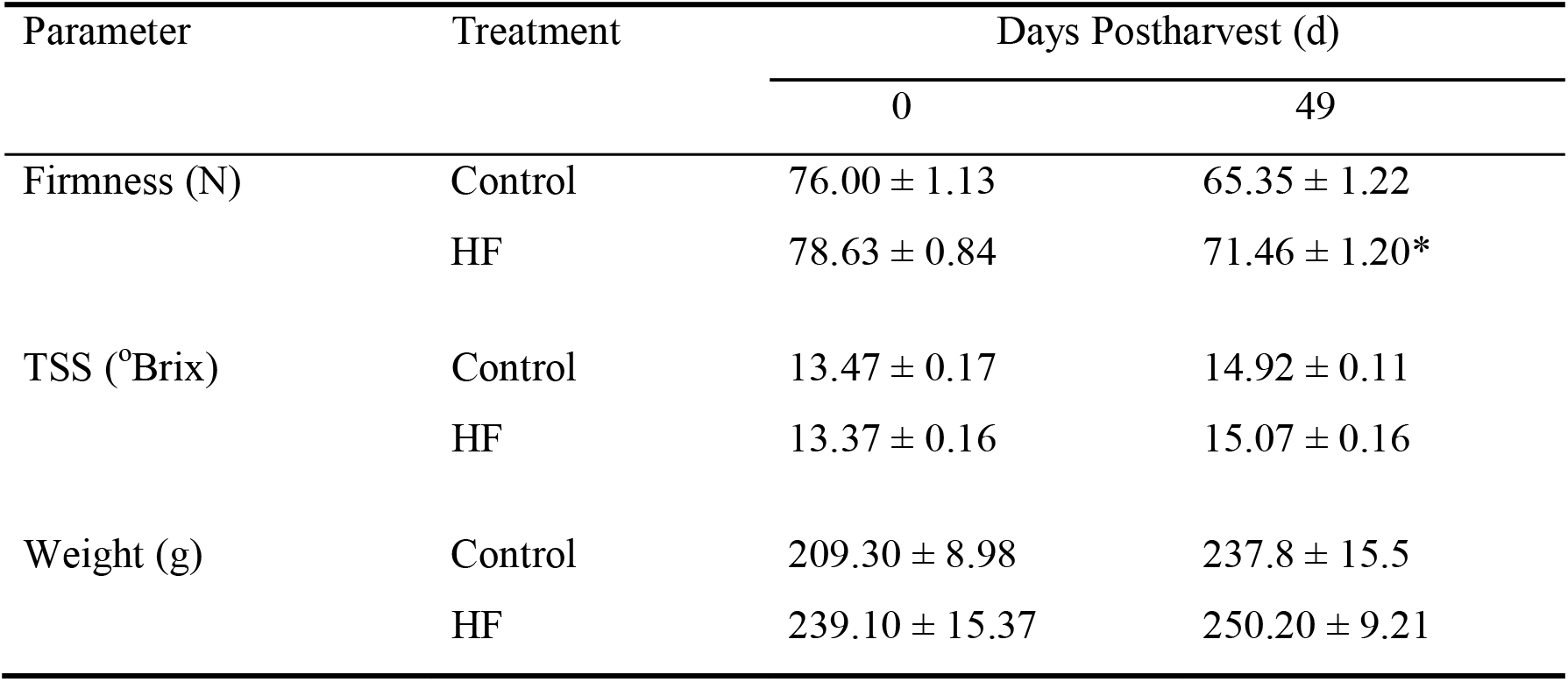
Fruit quality parameters of control and hexanal formulation treated ‘Honeycrisp’ apples at commercial harvest and 49 days after harvest. Values represent the mean ± SE of 10 randomly selected fruits. Means followed by asterisks indicate significant differences between control and hexanal formulation treatment at the same sampling time (*p* ≤ 0.05).

| Parameter | Treatment | Days Postharvest (d) |  |
| --- | --- | --- | --- |
|  |  | 0 | 49 |
| Firmness (N) | Control | 76.00 ± 1.13 | 65.35 ± 1.22 |
| | HF | 78.63 ± 0.84 | 71.46 ± 1.20* |
| TSS ($^{\circ}$Brix) | Control | 13.47 ± 0.17 | 14.92 ± 0.11 |
| | HF | 13.37 ± 0.16 | 15.07 ± 0.16 |
| Weight (g) | Control | 209.30 ± 8.98 | 237.8 ± 15.5 |
| | HF | 239.10 ± 15.37 | 250.20 ± 9.21 |

**Fig. 2:**
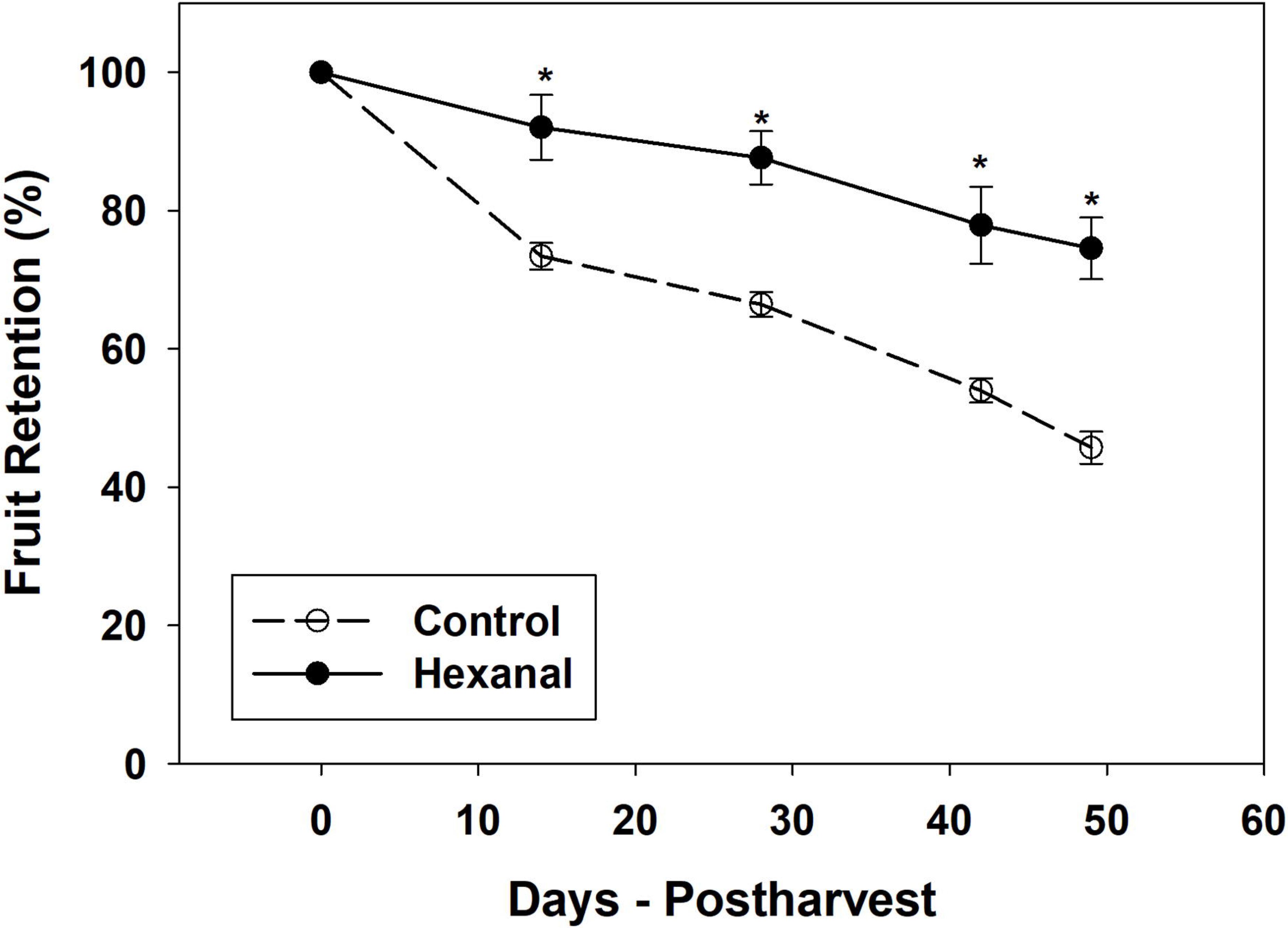
Percentage of Fruit retention in control and hexanal treated ‘Honeycrisp’ trees throughout the 49 days postharvest. Each value represents the mean ± SE of 4 trees. Asterisks indicate significant differences between control and hexanal treatment at the same sampling time (*p* ≤ 0.05).

### Quantitation of plant hormones in fruit and FAZ

At the commercial ripening stage, hexanal treated fruits produced lower ethylene (21%) than the control fruits (Table 2). Although it is not statistically different (*p* ≤ 0.05), lower ethylene at the commercial ripening stage obviously slows down the fruit ripening. Further, hexanal treatment significantly reduced both abscisic acid (ABA) and melatonin concentrations in the FAZ. However, hexanal did not alter the zeatin concentration in the FAZ (Table **2**). We could not detect any other plant hormones such as indole-3-acetic acid (IAA), gibberellic acid (GA3), salicylic acid (SA) and jasmonic acid (JA) in the FAZ samples. Presumably, those are present below the detection limit of the UPLC-MS (Waters, Canada).

**Table 2:** Variation of plant hormones in control and hexanal formulation treated ‘Honeycrisp’ fruit and fruit abscission zone (FAZ). Ethylene was measured in randomly selected eight fruits per treatment, and other plant hormones represent the mean ± SE of 18 replications of FAZ tissues harvested from two commercial orchards. Means followed by asterisks indicate significant differences between control and hexanal treatment (*p* ≤ 0.05).

| **Treatment** | **Ethylene**<br>($\mu\text{L L}^{-1} \cdot \text{kg}^{-1} \cdot \text{h}^{-1}$) | **Absciscic acid**<br>(ng. $\text{g}^{-1}$, DW) | **Zeatin**<br>(ng. $\text{g}^{-1}$, DW) | **Melatonin**<br>(ng. $\text{g}^{-1}$, DW) |
| --- | --- | --- | --- | --- |
| Control | $5.38 \pm 0.88$ | $320.17 \pm 33.25$ | $465.23 \pm 45.65$ | $56.58 \pm 6.37$ |
| Hexanal | $4.24 \pm 0.53$ | $192.99 \pm 11.83^*$ | $441.03 \pm 16.53$ | $39.81 \pm 2.47^*$ |

### Identification of differentially expressed genes

Mapping the rRNA depleted 96.89 million RNA-seq reads from the four samples against the apple reference genome (*Malus domestica*, GDDH13 v1.1.) showed that 92.99 million reads (96%) were mapped in total. The mean mapped reads per sample were 23.25 ± 0.28 million. After removing low expressed genes with less than ten raw reads across all samples, enabled the identification of 30,709 genes at least in one of the four samples (Table S2). EdgeR modelling following the ComBat batch correction and TMM normalization, yielded 726 DEGs between hexanal treated and control samples at *p* ≤ 0.05 with an expression foldchange cut off (FC) ≥ 2. Among the 726 DEGs, 353 were up-regulated (*p* ≤ 0.05; |log_2_foldchange ≥1|) while 373 were down-regulated (*p* ≤ 0.05; |log_2_foldchange ≥−1|) (Fig. 3A, B, Table S3) by hexanal.

**Fig. 3:**
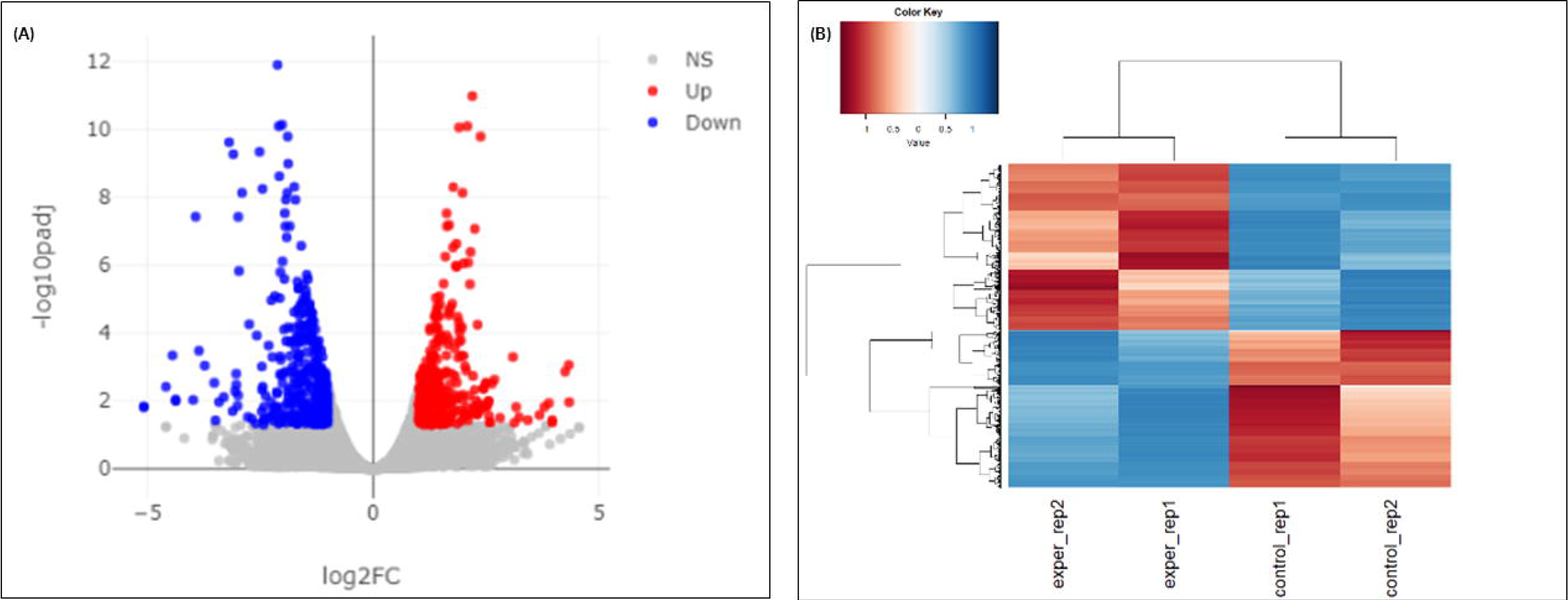
(A) represents the volcano plot of differentially expressed genes (DEGs). Red and blue dots indicate up (353) and down-regulated (373) genes and ash dots represent the nonsignificant genes in the FAZ. (B) represents heat map of DEGs in four samples. Blue and red show the down and up-regulated gene expression due to hexanal application, respectively. Additional information about the DEGs is presented in supplementary data Table S3.

### Identification of enriched gene ontology (GO) and functional pathways

Enrichment analysis was conducted to identify the enriched gene ontology (GO) functional classes and pathways related to the DEGs. We used *Arabidopsis thaliana* ortholog genes of the *Malus domestica* DEGs for this analysis. A total of 659 *Arabidopsis thaliana* ortholog genes were retrieved from the 726 *Malus domestica* DEGs. Enriched functional pathways were identified under all three GO term functional classes, biological process (BP), cellular component (CC) and molecular function (MF). The highest number of enriched pathways were identified under BP (231), while the lowest was under CC (9) functional classes (Fig. 4, Table S4). Hierarchical clustering and interactive network plot maps showed the correlation and relationship among the significant pathways, respectively (Fig. S1). Although various pathways could be identified under each GO term class, we focused on specific pathways related to fruit retention. These pathways were grouped into three main categories: plant hormone responses, transcription factors, and cell-wall modification. The DEGs belonged to selected enriched functional pathways were characterized.

**Fig. 4:**
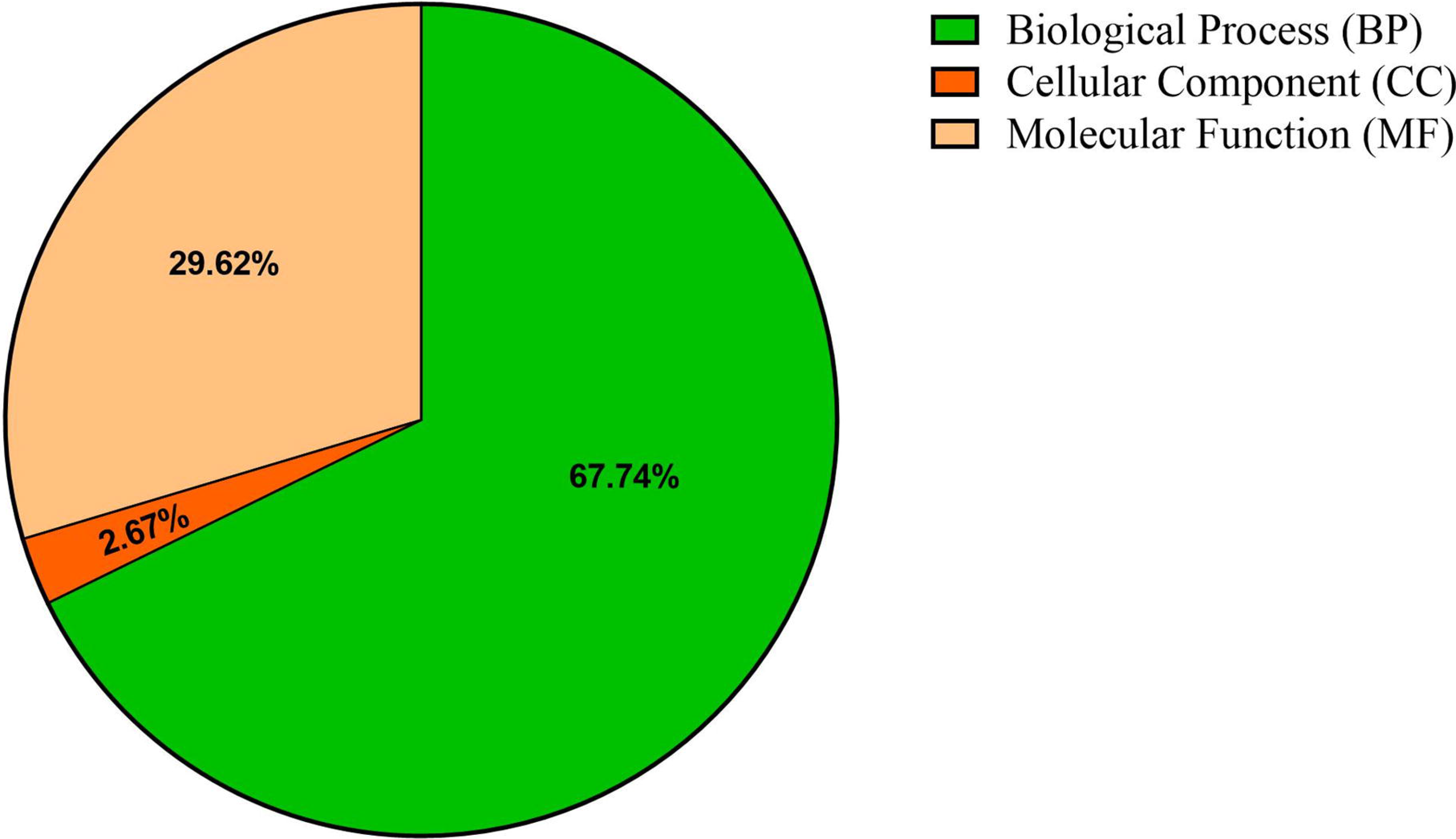
Number of enriched functional categories belonged to each Gene Ontology classification such as biological process (BP), cellular component (CC) and molecular function (MF). Hexanal altered greater percentage of BP (68%) and lesser of cellular components (3%). Enrichment was identified using Benjamini-Hochberg correction with a false discovery rate (FDR) at *p* ≤ 0.05. Additional information about the functional pathways is presented in supplementary data, Table S4.

### Characterization of genes related to various plant hormone responses

A total of sixty one DEGs related to various plant hormone responses were identified. Of these, genes related to ethylene (16), ABA (16), auxin (9) were the most represented, followed by JA (6), GA (5), SA (4 genes), cytokinin (4) and BR (1) showed the significant expression changes in response to hexanal (Table S5). Four genes involved in the ethylene biosynthesis pathway were identified, and all were down-regulated by hexanal, including S-adenosylmethionine synthase 2 (*SAM2*) and three, 1-aminocyclopropane-1carboxylic acid oxidase (*ACO3*, *ACO4* and *ACO4*-*like*). Two classes of ethylene receptors (*ETR2-like, ERS1*) were also identified, and both were up-regulated by hexanal. Ethylene signalling pathway elements were differentially expressed. Some transcriptional activators *AP2*/*ERF023*, *MYB113*-*like* were down-regulated, while others (*AP2*/*ERF017*, *AP2*/*ERF5*, *AP2*/*EREBP6*, *AP2*/*EREBP105* and *AP2/ERF*-*B3*-*RAV1*) were up-regulated by hexanal (Fig. 5A).

**Fig. 5:**
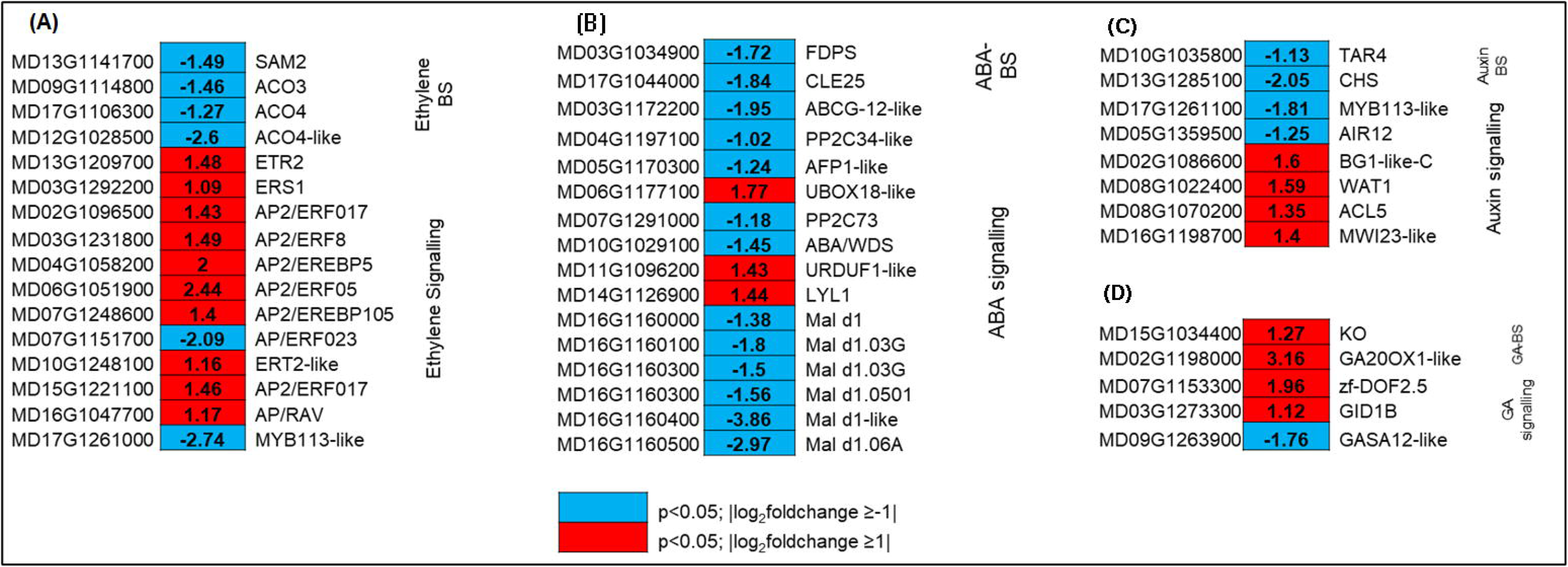
Expression profiling of genes related to biosynthesis and signalling of plant hormones (A) ethylene, (B) ABA, (C) auxin, and (D) GA. Blue and red represent down-and up-regulated gene expression due to hexanal application in the FAZ at harvest. The left column shows the *Malus domestica* gene id, the middle column shows the gene expression with |log_2_foldchange| values, and the right column shows the corresponding gene id. Additional information on the hormone-related genes is presented in supplementary data, Table S5.

Out of sixteen ABA-related DEGs, hexanal up- and down-regulated four and twelve genes, respectively (Fig. 5A). Two identified genes related to ABA biosynthesis, farnesyl diphosphate synthase (*FDPS*) and CLAVATA3-related protein-25 (*CLE25*), were down-regulated. Hexanal also down-regulated ten genes related to ABA signalling, including serine/threonine phosphatases PP2C (*PP2C*-*34*-*like*, *PP2C*-*73*), ninja-family protein-*AFP1*-*like*, and ABC transporter G family member, *ABCG12*-*like*. On the contrary, most of the GA related genes were up-regulated. The key genes involved in GA biosynthesis such as gibberellin 20 oxidase1-like (*GA20OX*-*like*) and ent-kaurene oxidase (*KO*) were up-regulated by nine and two fold, respectively by hexanal. Besides, a receptor gene *GIDIB* was up-regulated by two fold compared to control (Fig. 5D). Gene expression showed a divergent pattern in auxin biosynthesis, signalling, transport, and response. Genes related to auxin biosynthesis, tryptophan aminotransferase-related protein 3 (*TAR3*) and response (*CHS*, *MYB113*-*like* and *CYP75B1*) were down-regulated by hexanal in the FAZ, while genes involved in auxin transports such as protein big grain 1-like (*BG1*), protein walls are thin-1 (*WAT1*), and thermospermine synthase-5 (*ACL5*) were up-regulated (Fig. 5C).

Altogether, fifteen DEGs were identified related to SA, JA, cytokinin, BR biosynthesis and signalling. The expression of these genes was greatly varied. For example, three out of four genes related to SA were up-regulated. Whereas three out of six genes related to JA were down-regulated with the expression fold change between two and seven. Genes related to cytokinin were mostly down-regulated (Table S5). The above results suggested eight classes of plant hormones were altered by hexanal at the commercial ripening stage.

### Characterization of genes encoding for transcription factors

A total of twenty one genes putatively encoding TFs of diverse families were differentially expressed in the FAZ (Table S6). Of those, most genes (8) belonged to APETALA2/ERF (AP2/ERF) superfamily. Seven genes representing AP2/ERF family were up-regulated while one was down-regulated by hexanal. These genes either act as repressor (*AP2/ERF4*) or activators (*AP2/ERF017, AP2/EREBP6, AP2/ERF5, AP2/ERF023*, *AP2/EREBP105 and AP2/ERF*-*B3*-*RAV1*) of GCC-box mediated gene expression in ethylene-activated signalling pathway. Besides, few genes belonged to TF families bHLH, GATA, MADS-box, MYB, TCP and WRKY were also identified. MYB113-like gene representing MYB transcription factor family, participating in anthocyanin biosynthetic process was down-regulated by hexanal at harvest.

### Characterization of genes related to cell-wall modification

A total of thirty-two genes related to cell-wall modifications were differentially expressed in the FAZ (Fig. 6, Table S7). Of these, twenty six and six were down-and up-regulated, respectively. The genes encoding enzymes related to callose, polygalacturonase and expansins were down-regulated by hexanal. Of those down-regulated genes, eight were related to callose degradation, including endoglucanase 19-like, endo 1,4-β-glucanase, endoglucanase-45-like and glucan endo-1,3 β-glucosidase 8-like. Two genes were related to polygalacturonase (PG), including polygalacturonase 1, endo-polygalacturonase-like-protein-like. Whereas seven genes related to expansin, including EXPA1-like, EXPA6, EXPA8, EXPA10-like, EXPA16-like (Fig. 6). All these genes encoding enzymes related to callose, PGs, and expansins could be involved in maintaining the cell-wall integrity of the FAZ cells

**Fig. 6:**
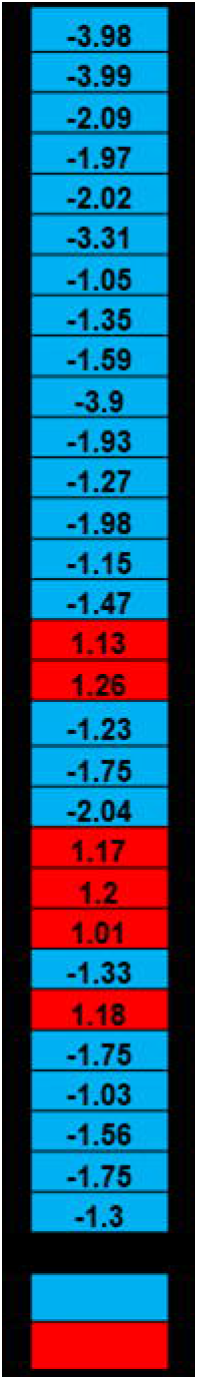
Expression profiling of genes related to cell-wall modification. Blue and red represent down-and up-regulated gene expression due to hexanal application in the FAZ at harvest. The left column shows the *Malus domestica* gene id, the middle column shows the gene expression with |log_2_foldchange| values, and the right column shows the corresponding gene id. Additional information on the hormone-related genes is presented in supplementary data, Table S7.

### Characterization genes related to abscission

Two genes specifically involved in abscission were identified. Senescence-associated carboxylesterase 101-like (*SAG101*-*like*) encodes an acyl hydrolase involved in senescence/floral organ abscission that was down-regulated. In contrast, zinc finger protein 2-like (*ZFP2*-*like*) that acts as a negative regulator of floral organ abscission was up-regulated by hexanal.

### Confirmation of gene expression patterns by qRT-PCR

All tested genes belonged to ethylene biosynthesis and signalling (*SAM2;* MD13G1141700, *ACO3;* MD09G1114800, *ETR2*-*like*; MD13G1209700 and *ERF17;* MD15G1221100) (Fig. 7A) and cell-wall metabolisms (*EXPA6*; MD03G1090700, *EXPA8*; MD07G1233100, *EG19-like;* MD06G1105900 and *1,4*-*β*-*EG3;* MD10G1003400) (Fig. 7B) were identified by the qRT-PCR analysis. Moreover, our qRT-PCR results revealed down-regulation of all eight selected genes related to ethylene biosynthesis and cell-wall modifications, which is identical to the RNA-Seq results. Thus, the qRT-PCR results were consistent with the RNA-seq data, confirming the RNA-seq analysis.

**Fig. 7:**
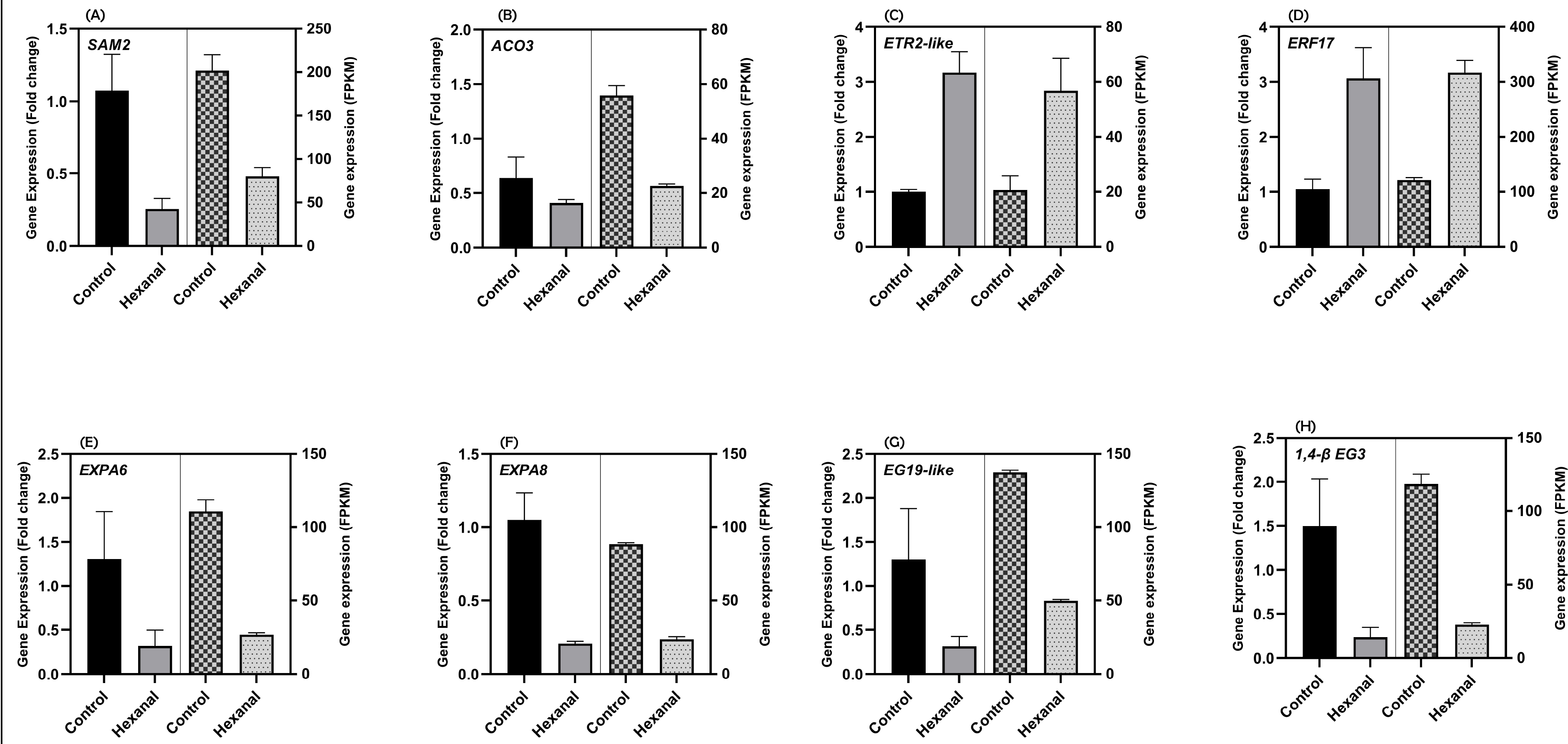
qRT-PCR confirmation of gene expression pattern of selected eight genes representing (A)-(D) ethylene biosynthesis and signalling and (E)-(H) cell-wall modification. The data represents the mean values of four biological replicates and three technical replicates representing the samples harvested from both commercial orchards. FPKM values of each gene were calculated from RNA-Seq reads counts normalized to a per million total reads counts. Genes and the primers are shown in the supplementary data, Table S1.

**Fig. 8:**
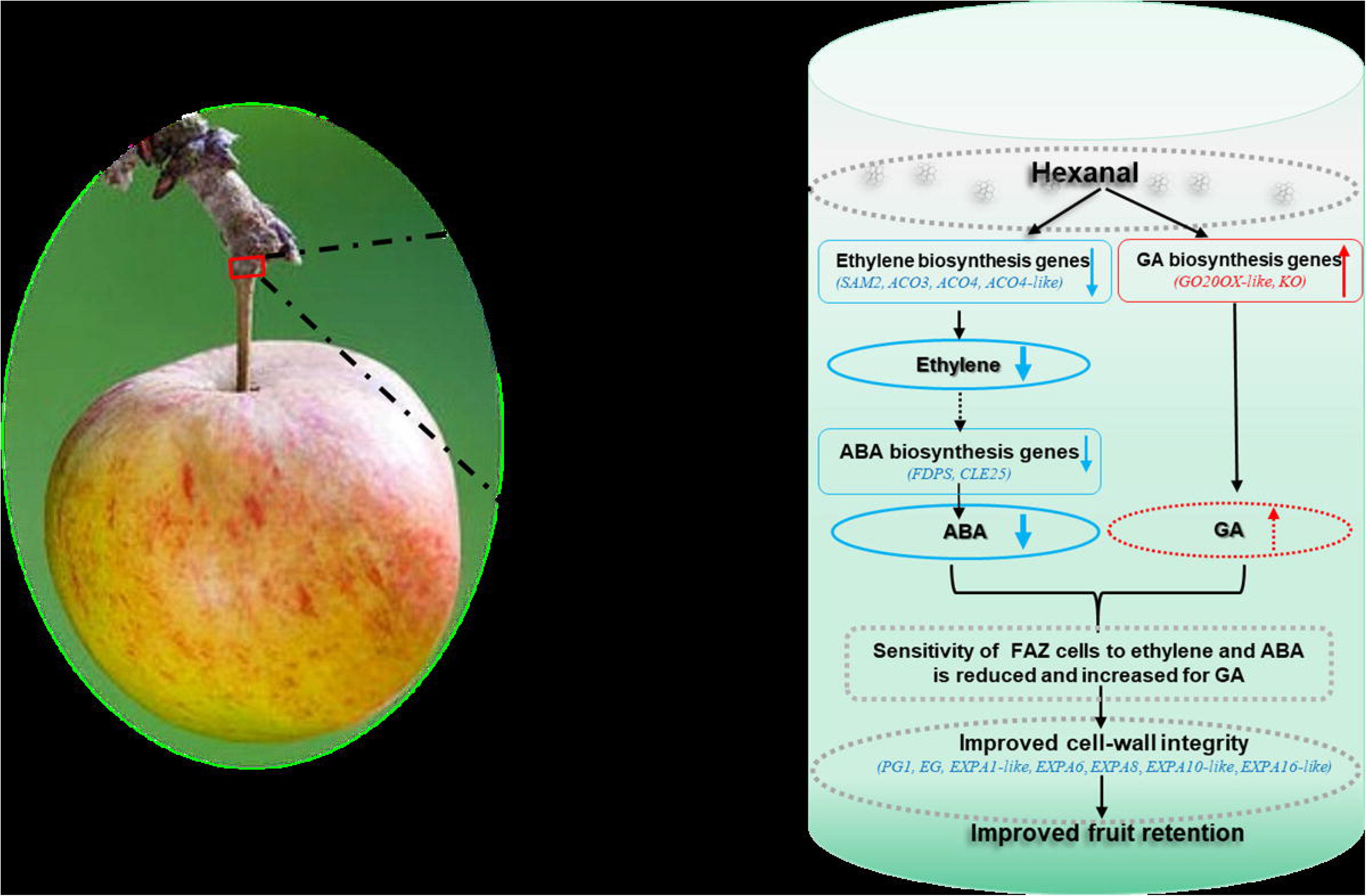
Proposed model of hexanal mediated fruit retention in ‘Honeycrisp’ apples. Preharvest hexanal spray downregulated the expression of genes involved in ethylene biosynthesis in the FAZ and thus decreased the ethylene. Lower ethylene, in turn slows down the expression of the ABA biosynthesis genes and substantially minimize the ABA level in the FAZ. At the same time, GA biosynthesis genes were up-regulated by hexanal and may enhance the GA concentration, which counters ABA. Hence, the sensitivity of FAZ cells to ABA decreased. Parallelly, hexanal also downregulated genes related to cell-wall degrading enzymes such as EG, PG, and expansins. Consequently, cell-wall integrity of the FAZ cells improved in the treated fruits. Collectively, these events improved the fruit retention of the hexanal treated fruits. “Solid arrows represent known mechanism; broken arrows represent unknown mechanism; blue represent down-regulation/decrease events, red represent up-regulation/increase events”.

## Discussion

### Ethylene production and fruit ripening are reduced in hexanal treated fruits

Fruit drop shortly before the harvest is a challenge to apple growers that causes significant yield losses. In climacteric fruits like apples, the production of ethylene by the ripening fruits stimulates the production of cell-wall degrading enzymes and forms an abscission zone in the pedicel (Taylor and Whitelaw, 2001). Growers extensively resort to multiple PGR applications to reduce ethylene production and slow down the ripening and improve fruit retention. However, the application of PGRs has its own limitations and further increases the cost of production (Li *et al*., 2010; Robinson *et al*., 2010). ‘Honeycrisp’ is a premium apple variety that fetches higher returns for the growers but is also more prone to pre-harvest fruit drop. So, reducing the pre-harvest drop in such premium varieties will boost the economic returns for the growers as well as reduce preventable food loss. The application of hexanal as a pre-harvest spray improved fruit retention in a several fruit crops (Anusuya *et al*., 2016; El Kayal *et al*., 2017). However, the detailed mechanisms on how hexanal improves fruit retention are not well understood. Here we attempted to elucidate the hexanal’s mode of action on improved fruit retention in ‘Honeycrisp’ by studying the hormonal variations, differential expression of genes and enriched functional pathways.

Hexanal treated fruits produced less ethylene and retained firmness for a longer time compared to control fruits (Table 1, 2). Our results are consistent with the previous observations of hexanal on similar climacteric fruits such as mango (Jincy *et al*., 2018) and banana (Yumbya *et al*., 2018). Moreover, hexanal spray had a striking effect on fruit retention for an extended time (Fig. 2). Extending fruit retention time would be valuable to growers as it can extend the harvesting window, reduce postharvest loss, thereby stabilizing the market price. Moreover, enhanced fruit firmness is an added advantage for premium apple variety like ‘Honeycrisp’ as they are mainly bred for the fresh market. The improvement in fruit retention, firmness and reduced ethylene are due to hexanal which slows down the ripening process, thus delaying the abscission.

### Hexanal interferes with ethylene biosynthesis and perception in the FAZ

Ethylene biosynthesis increases before abscission in many senescing organs, including fruits (Bonghi *et al*., 2000). In apple fruitlet, application of chemical thinner ethephon stimulated the ethylene biosynthesis in parallel with the upregulation of key regulatory genes, *MdACO1, MdACS5A* and *MdACS5B* in the FAZ (Kolarič *et al*., 2011) suggesting that, ethylene biosynthesis and signalling is involved in abscission. We have identified four key genes involved in the ethylene biosynthesis pathway, including *SAM2*, *ACO3*, *ACO4*, and *ACO4*-*like* (Fig. 5A, Table S5). Interestingly, the expression of all four genes was decreased by hexanal. ‘Red Delicious’ apples sprayed with ethylene suppressors AVG and 1-MCP resulted in decreased expression of *MdACS5A* and *MdACO1* in the FAZ and concomitant decreased fruit ethylene and fruit drop (Li and Yuan, 2008).

An important aspect of ethylene action in abscission is its perception and tissue sensitivity (Bonghi *et al*., 2000; Taylor and Whitelaw, 2001; Botton and Ruperti, 2019). Ethylene receptors serve as negative regulators to regulated ethylene response, and there is an inverse relationship between receptor levels and ethylene sensitivity of a tissue (Hua and Meyerowitz, 1998). The present results show that an increase in transcript levels in ethylene receptors *ERS1* and *ETR2*-*like* in response to hexanal in the FAZ at harvest (Fig. 5A; Table S5). Expression of both receptors could be due to the compensatory mechanism existing within the complex of ethylene receptors. A similar result was observed in AVG-treated nectarine, where *PpETR1* and *PpERS1* transcripts were overexpressed at harvest (Ziosi *et al*., 2006). Moreover, an increased expression of *Pp*-*ERS1* observed 1-MCP treated peach (Rasori *et al*., 2002), muskmelon (Lashbrook *et al*., 1998) and tomato (Sato-Nara *et al*., 1999). Likewise, a *LeETR4* was overexpressed in the *Never-ripe* (*NR*) antisense tomato besides the expected repression of *NR* transcript (Tieman *et al*., 2001). Together, the gene expression in the FAZ and concomitant decrease in ethylene production in fruits prove that fruit retention in ‘Honeycrisp’ by hexanal is likely to be ethylene dependent.

### Hexanal mediates hormonal crosstalk in the FAZ

Other plant hormones, including ABA, have a stimulatory effect in abscission (Zhu *et al*., 2011). However, whether the stimulatory effect is due to the direct involvement of ABA or mediated by the production of ethylene is still unclear. At commercial harvest, a parallel reduction in ABA level and expression of ABA biosynthetic genes *FDPS* and *CLE25* were observed in the hexanal treated FAZ (Fig. 5B; Table S5). Likewise, ABA signalling components *PP2C*-*34*-*like*, *PP2C*-*37*, and ABA response proteins, including allergen-like proteins (*Mal d1*, *Mal d1.03G*, *Mal d1*-*like* and *Mald.06A*) showed decreased expression. Earlier studies in melon mature fruit abscission revealed an up-regulation of *SnRK2*/PP2Cs that was attributed to early fruit abscission (Corbacho *et al*. 2013) suggesting that regulation of abscission related signalling compound trigger the onset of fruit drop and controlling them could lead to preventing such drops.

We can clearly observe a concomitant reduction in abscission promoting hormones ethylene and ABA by hexanal at harvest, suggesting that hexanal mediates a cross-talk between these hormones. In general, ABA and gibberellins (GA) are one pair of classic phytohormones, which antagonistically mediate several plant developmental processes including, fruit-abscission (Chen *et al*., 2008; Liu and Hou 2018). Interestingly, all the DEGs related to GA biosynthesis and signalling were up-regulated (Fig. 5D; Table S5), suggesting hexanal also mediates hormonal cross-talk between ABA and GA.

### Hexanal suppresses genes associated with cell wall degradation and abscission

The abscission starts with the expression of several wall-loosening enzymes such as cellulases, polygalacturonases and expansins. The collective action of all these enzymes accelerates the dissolution of the middle lamella, resulting in organ separation (Bonghi *et al*., 1993; Li *et al*., 2010, Merelo *et al*., 2017). Enlargement of AZ cells involves cell wall loosening, which can be aided by expansins (Choe and Cosgrove, 2010). Several reports have reported that expansins are expressed abundantly in AZ including tomato flower (Tsuchiya *et al*., 2015) and apple fruit (Zhu *et al*., 2011). The present study has identified seven genes encoding expansins (*EXPs*), showed decreased expression due to hexanal (Fig. 6; Table S7). Certain AZ cells enlarge in response to ethylene (Osborne, 1976). Microscopic visualization of hexanal treated FAZ cells were smaller, more organized with more defined horizontal layers than control cells (Fig. S2). Hexanal presumably reduced ethylene-mediated abscission by the suppression of expansins.

Our results showed that *MdPG1*expression was decreased by hexanal in the FAZ (Fig. 6; Table S7). Increase in polygalacturonase (PG) activity correlates with fruit abscission (Tonuti *et al*., 1995; Bonghi *et al*., 2000). *MdPG1* was involved in apple fruit softening, whose expression was reduced by 1-MCP and AVG treatments (Li and Yuan, 2008). Increase in endo-β-1,4-glucanase (EG) activity has been related to fruit abscission in several crops, including apple (Bonghi *et al*., 2000). Interestingly, all identified EGs from RNA-seq and qPCR analysis were down-regulated by hexanal (Fig. 6; Table S7). Moreover, decreased expression of an abscission specific gene, *SAG101*-*like* (MD17G1039700), encodes an acyl hydrolase involved in senescence/floral organ abscission may assist in retaining the fruits in hexanal treated trees.

In conclusion, this work demonstrates the crucial role of hexanal in improving fruit retention and fruit qualities. The mechanism of improved fruit retention by hexanal in ‘Honeycrisp’ is likely through an ethylene dependent pathway. Hexanal decreased the ethylene production in fruits, thus may reduce the sensitivity of FAZ cells to ethylene and ABA through down-regulating the genes associated with ethylene and ABA biosynthesis in the FAZ. Besides, hexanal can maintain the cell-wall integrity of abscission zone cells by modulating cell-wall degrading enzymes such as expansins, EGs and PG. Thus, hexanal application promises to be a great technology to control fruit drop in ‘Honeycrisp’ apples, given that this cultivar is categorized as more prone to fruit drop. Hexanal is a natural compound produced by all ripening fruits and generally regarded as a safe compound (GRAS). Also, it has been approved by FDA as a food additive. Further studies could be directed to validate how hexanal slows down the ethylene signal from fruit to the AZ.

## Supplementary data

Supplementary Fig.S1: (A) interactive plots and (B) Hierarchical clustering of GO term biological process

Supplementary Fig S2: Microscopic visualization of fruit-AZ in (A) control and (B) hexanal treatment

Supplementary Table S1: List of primers used for the qRT-PCR analysis Supplementary Table S2: Summary results of RNA-seq data

Supplementary Table S3: List of differentially expressed genes

Supplementary Table S4: Enriched GO terms and functional pathways

Supplementary Table S5: DEGs related to plant hormones

Supplementary Table S6: DEGs related to transcription factors

Supplementary Table S7: DEGs related to cell-wall modification

## Acknowledgment

Financial support for this project was provided by Canada’s International Development Research Centre (IDRC) by the Government of Canada through Global Affairs Canada (GAC), for JS. The Arrell Food Institute at the University of Guelph, Margaret and Angus Hamilton Apple Tree Fruit Research scholarship, Keith and June Laver scholarship in Horticulture and Vineland Centennial Horticultural scholarship at the University of Guelph to KS. We gratefully acknowledge the technical support of Glen Alm (Senior Research Technician) and the growers, Feenstra and family, and Art and Marlene Moyer for providing their orchards for experimentation. We would also like to thank Erika DeBrouwer (Tree Fruit Specialist, OMAFRA), for her support on this project.

## Abbreviations

ABA: abscisic acid
ACO: 1-aminocyclopropane-1carboxylic acid oxidases
DEGs: differentially expressed genes
EG: endo-glucanases
EXP: expansins
FAZ: fruit abscission zone
GA: gibberellic acid
PGRs: plant growth regulators
PG: polygalacturonase
PFD: preharvest fruit drop
RNA-seq: RNA sequencing
SAM: S-adenosylmethionine synthase

## Author Contributions

KS conducted the experiments, analysed the data and wrote the manuscript, WE helped in running and analysing the qRT-PCR results, DT helped in analysing the transcriptomic data, MMA and PKS helped in running and interpreting the hormone analysis results, AS and GP helped in the conceptualization and planning of the experiment and JS conceptualized the experiments, supervised and edited the manuscript.

## Data Availability Statement

All data supporting the findings of this study are available within the paper and within its supplementary materials published online.

